# Faunal and Floral Assemblages of the Zavitan (Zvitan) Stream near Katzrin (Central Golan Heights, Israel) and Their Influence on Amphibians

**DOI:** 10.64898/2026.04.26.718926

**Authors:** Monika Almozlino, Gad Degani, Dani Bercovich, Ari Meerson

## Abstract

The Zvitan (Zavitan) Stream is one of the major basaltic drainage systems of the Levant Region in Israel. The present short communication focuses specifically on the sector adjacent to Katzrin, integrating geomorphological, hydrological, ecological, and amphibian distribution data within the broader watershed context. The stream originates in the central Golan plateau and flows westward into the Yehudiya Reserve before joining the Meshushim Stream and ultimately draining into Lake Kinneret. In the Katzrin sector, the stream is characterized by deeply incised basalt canyons, winter spring discharge and semi-permanent pools sustained by springs and seepage. Seasonal hydrological fluctuations strongly influence aquatic habitats and amphibian breeding success. Amphibian species documented in the Golan Heights include *Salamandra salamandra infraimmaculata, Triturus vittatus* (currently *Ommatotriton vittatus*), *Hyla savignyi, Bufotes viridis, Pelophylax bedriagae* (formerly *Rana ridibunda*), and *Pelobates syriacus*. Their distribution is closely associated with water availability, elevation, temperature, and hydroperiod, as demonstrated in northern Israel habitats. The Katzrin sector of Zvitan represents an intermediate ecological zone where Mediterranean and steppe elements converge, creating heterogeneous amphibian assemblages. This observation, carried out at a specific site in the Zavitan Stream, aimed to examine the ecological conditions and identify which amphibian species inhabit this pool, where environmental conditions may differ from those in the main stream channel.

## Introduction

The Zavitan Stream (also spelled Zvitan or Zavit) is one of the principal tributaries of the Yehudiya drainage system in the central Levant Region (Syria–Lebanon–Israel). Syria, Lebanon, and Israel belong primarily to the Levantine subregion of the Eastern Mediterranean biogeographical province, which forms part of the Palearctic Realm (Cox and Moore, 2001; Udvardy, 1975). In classical zoogeographical terminology, this area is often referred to as the Syro-Levantine region or Levantine corridor, a transitional zone between the Mediterranean, Irano-Turanian, and Saharo-Arabian biogeographic regions (Por, 1975; Yom-Tov and Tchernov, 1988).The term Levant Corridor is especially important in zoology and paleo-biogeography, as it describes the main dispersal route between Africa and Eurasia during the Neogene and Quaternary periods (Tchernov, 1988). The fauna of Syria, Lebanon, and Israel belongs primarily to the Levantine subregion of the Eastern Mediterranean province within the Palearctic Realm. This area represents a major biogeographical transition zone and dispersal corridor between Afro-tropical, Irano-Turanian, and Mediterranean faunal elements (Por, 1975; Tchernov, 1988; Cox, 2001; Roll et al., 2009).

The stream drains a basaltic plateau formed by extensive Miocene–Pliocene volcanic activity and flows westward toward the upper Jordan River basin. The sector located near Katzrin represents a geomorphologically and ecologically distinctive segment characterized by deeply incised canyons, perennial spring inputs, seasonal runoff pulses, and a mosaic of aquatic and riparian habitats. The geomorphology of the Zavitan basin is controlled primarily by basaltic bedrock, tectonic uplift, and fluvial incision processes. The hard basalt substrate promotes waterfall formation, plunge pools, and narrow canyon morphology, while joints and fractures facilitate groundwater recharge and spring emergence (Sneh and Weinberger, 2003; Gvirtzman, 2009). The hydrological regime of the stream is Mediterranean, with most discharge occurring during winter precipitation events (November–March), followed by substantial reduction in flow during the dry summer season. However, localized perennial flow is maintained in certain reaches due to spring discharge and groundwater seepage, particularly within shaded canyon sections (Gafni et al., 2000; Zohary and Ostrovsky, 2011).

Water quality in northern Israeli basaltic streams is generally characterized by moderate conductivity, circumneutral to slightly alkaline pH, and relatively low nutrient concentrations compared to lowland agricultural streams (Gafny et al., 2000; Zohary and Ostrovsky, 2011). Nevertheless, seasonal fluctuations in temperature, dissolved oxygen, and nutrient availability occur due to hydrological variability and increasing anthropogenic pressures, including tourism and land-use change in the Katzrin region. Such limnological variability directly influences aquatic community structure, especially macroinvertebrates, fish, and amphibians.

Riparian vegetation along the Zavitan sector near Katzrin reflects the transition between open basalt plateau vegetation and canyon-confined Mediterranean woodland. The plateau is typically dominated by Mount Tabor oak (*Quercus ithaburensis*) associations and shrubland elements, while the canyon bottom supports hygrophilous vegetation including *Nerium oleander, Salix* spp. and *Phragmites australis* in wetter microsites (Zohary, 1973; Danin, 2004). This structural heterogeneity provides critical habitat for amphibians and other vertebrates.

Amphibian assemblages in the central Levant Region are shaped by hydroperiod, water permanence, temperature regime, and habitat connectivity (Gafny et al., 2000; Preißler, 2021). Streams and associated rock pools within the Zavitan drainage may support species such as the Levant water frog (*Pelophylax bedriagae*), the Near Eastern fire salamander (*Salamandra infraimmaculata*) and the green toad complex (*Bufotes viridis* group), depending on hydro ecological conditions and seasonal water persistence. The canyon pools serve as breeding sites or larval development habitats (Degani and Kaplan, 1999), while adjacent terrestrial habitats provide refugia during dry periods. Because Mediterranean streams are highly sensitive to climatic variability, amphibian populations in these systems may serve as biological indicators of hydrological stability and ecosystem integrity (Degani, 2019, 2024b).

The Katzrin–Zavitan sector is also embedded within the broader Yehudiya Nature Reserve landscape, an area of high conservation value that supports diverse reptile, bird, and mammalian communities. Increasing recreational use, climate warming trends, and altered precipitation regimes may influence stream discharge patterns, pool permanence, and amphibian reproductive success in the coming decades (Zohary and Ostrovsky, 2011). Therefore, a synthesis of geological, hydrological, limnological, and biological information specific to the Zavitan stream sector near Katzrin is essential for understanding ecosystem function and guiding conservation management.

The aim of the present observation was to investigate the ecological conditions, with particular emphasis on water temperature, in a small water pool located along the Zavitan Stream near Katzrin (Central Golan Heights, Israel). The study also sought to document the amphibian species present in this pool and to compare their occurrence with the amphibian community known from the surrounding region.

## Materials and Methods

### Study Site

The study was conducted at a seasonal freshwater pool located near the village of Katzrin in the central Golan Heights, northern Israel. The pool (hereafter “Katzrin Pool”) is situated at 35°42′25″N, 35°59′12″E (altitude: 339 m above sea level) (Figures 1–2). The site lies within a basaltic plateau landscape characteristic of the central Golan, where winter rainfall accumulates in shallow depressions, forming temporary lentic habitats (Degani and Kaplan, 1999).

**Figure 1.**
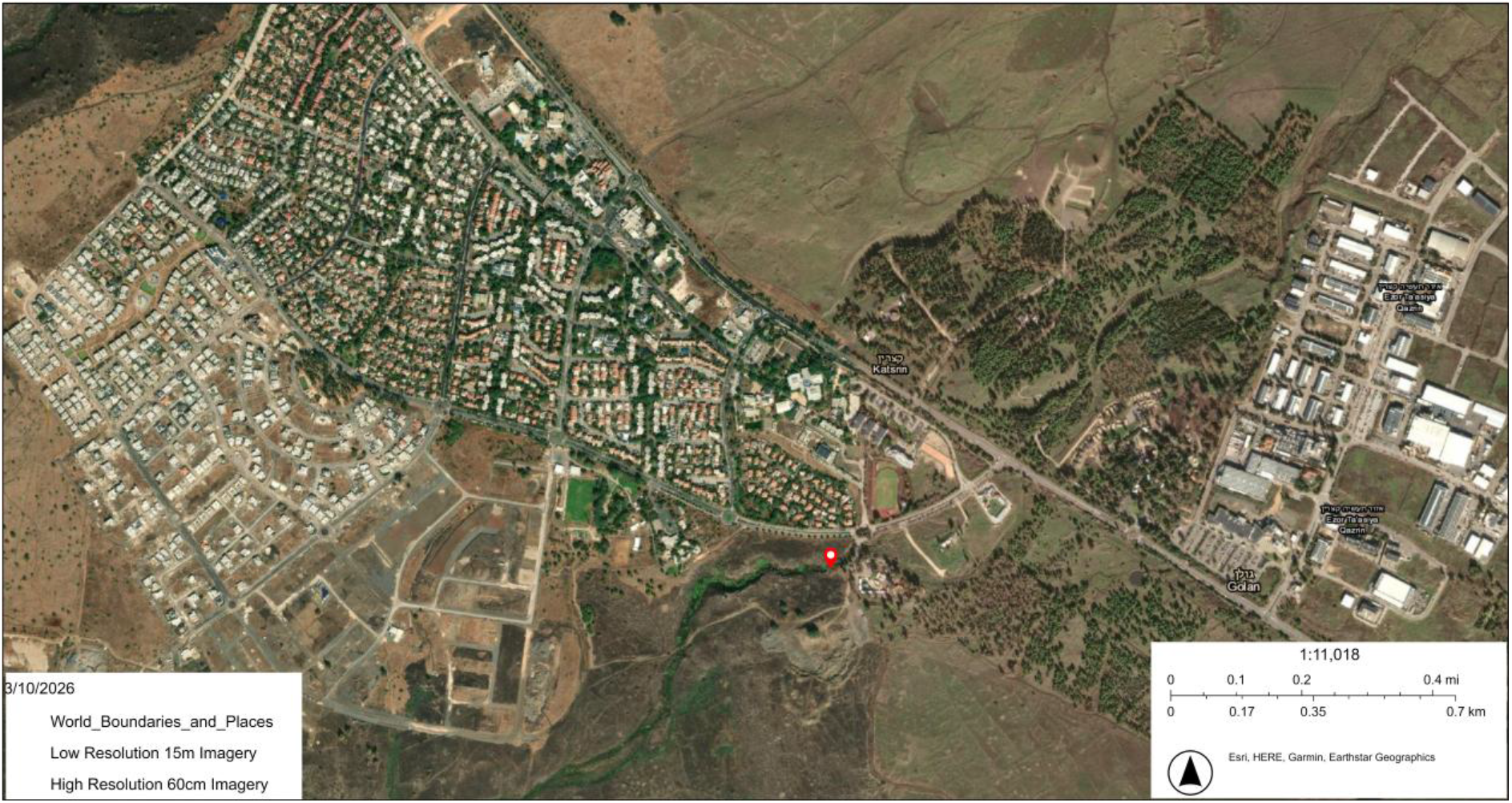
Location of the study site in the Zavitan Stream near Katzrin, with the exact observation point indicated by a red dot (base map: satellite imagery with labels, moag.maps; https://moag.maps.arcgis.com/).

**Figure 2.**
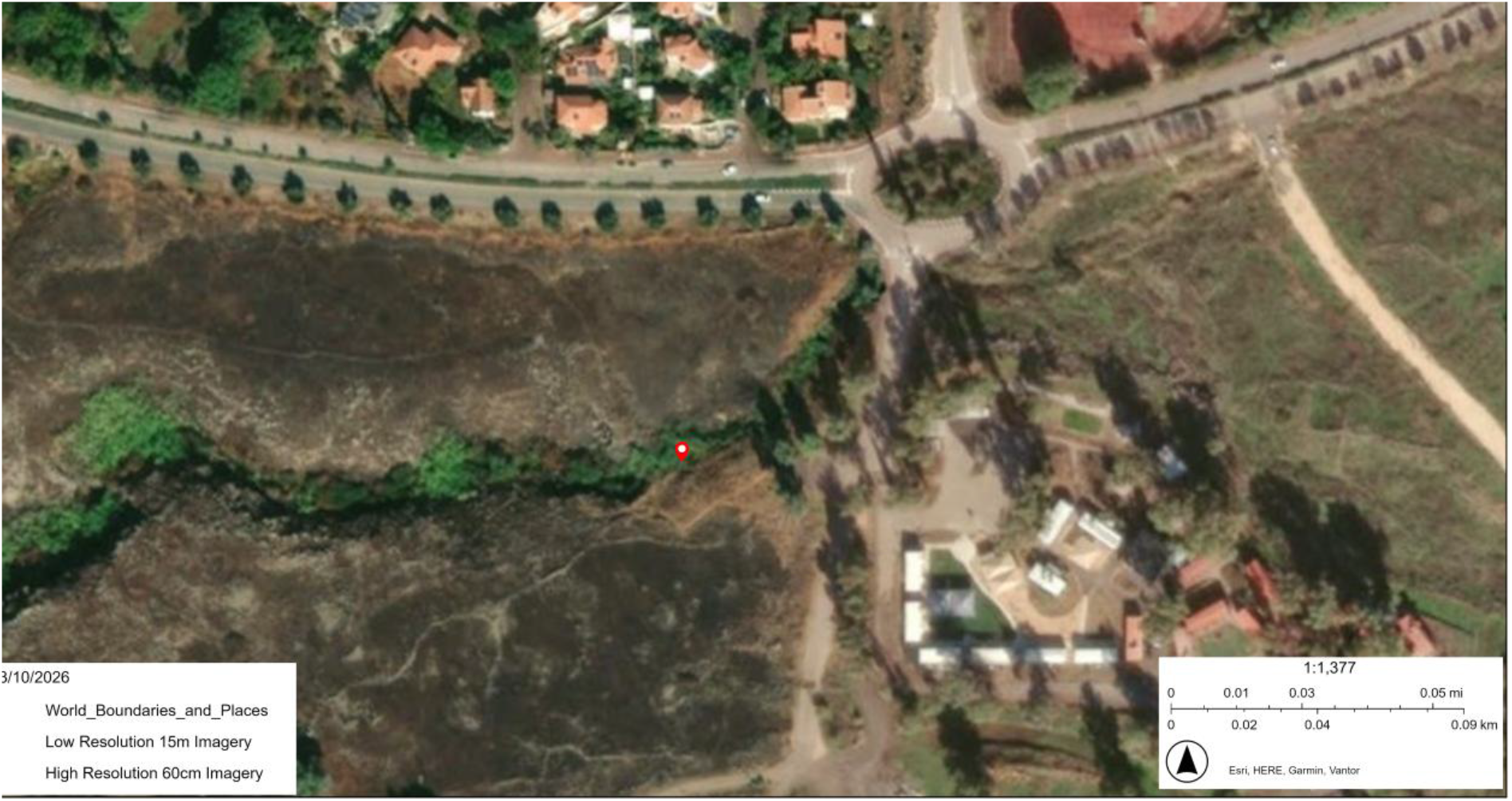
Close-up view of the Katzrin pool in the Zavitan Stream near Katzrin, with the exact observation site indicated by a red arrow (base map: satellite imagery with labels, Moag.maps).

Katzrin Pool is rain-fed and seasonal, typically filling during winter precipitation events (December–January) and gradually drying during late spring and early summer (June–July). The hydroperiod length is primarily determined by annual rainfall, evaporation rates, and substrate permeability. The pool provides temporary breeding habitat for amphibians and supports associated aquatic invertebrates and emergent vegetation during the inundation period.

### Monitoring Schedule

Field monitoring was conducted from January to July 2025, encompassing the complete hydroperiod of the pool and the known breeding season of the amphibians (Degani and Kaplan, 1999). During the initial inundation phase (January–February), the pool was monitored weekly in order to document early colonization and breeding activity. Subsequently, monitoring was conducted on a monthly basis until complete desiccation of the pool in July 2025. The termination of monitoring coincided with total water loss and the end of the amphibian reproductive period (Degani and Kaplan, 1999).

### Environmental Data Collection

During each field survey, water temperature (°C) was measured *in situ* using a calibrated digital thermometer at a depth of approximately 10 cm below the water surface. The hydrological status of the pool was qualitatively assessed and categorized according to water level conditions (e.g., full, declining, shallow, isolated puddles, or dry). In addition, local meteorological conditions, including estimated air temperature, were recorded.

### Biological Sampling

Aquatic organisms were sampled using a hand net (Pawfly Aquarium Fish Net; frame size 10″ × 7″; pocket depth 5″) (Degani and Goldberg, 2013 ; Goldberg et al., 2009; Goldberg et al., 2012; Pearlson and Degani, 2007) and transferred to a transparent plastic container for examination. Sampling effort during each visit included sweeping the net through shallow vegetated margins, sampling open-water areas and inspecting submerged vegetation and detritus.

Captured organisms were identified to the lowest possible taxonomic level in the field. Particular attention was given to amphibians. After identification and documentation, all individuals were immediately returned to the pool. Additionally, all plant and animal species observed at the site (aquatic and riparian) were recorded through direct observation (Degani, 2024a; Degani and Ahkked, 2021; Pearlson and Degani, 2007). All fieldwork was conducted under permit no. 2024-43687 issued by the Israel Nature and Parks Authority.

## Results

### Abiotic Observations

A total of nine monitoring visits were conducted between January and July 2025, encompassing the full hydroperiod of the seasonal pool. The pool retained standing water throughout the monitoring period until July 2025, when complete desiccation was recorded. No residual puddles remained after this date, indicating the termination of the aquatic phase for the 2025 season.

Water temperature (Figure 3) exhibited moderate seasonal variation over the study period. Recorded values ranged from 14.8 °C (early winter monitoring) to 21.3 °C (late spring), with a mean water temperature of 18.49 °C. A gradual increase in water temperature was observed from January toward late spring, corresponding with rising ambient temperatures and progressive reduction in water depth.

**Figure 3.**
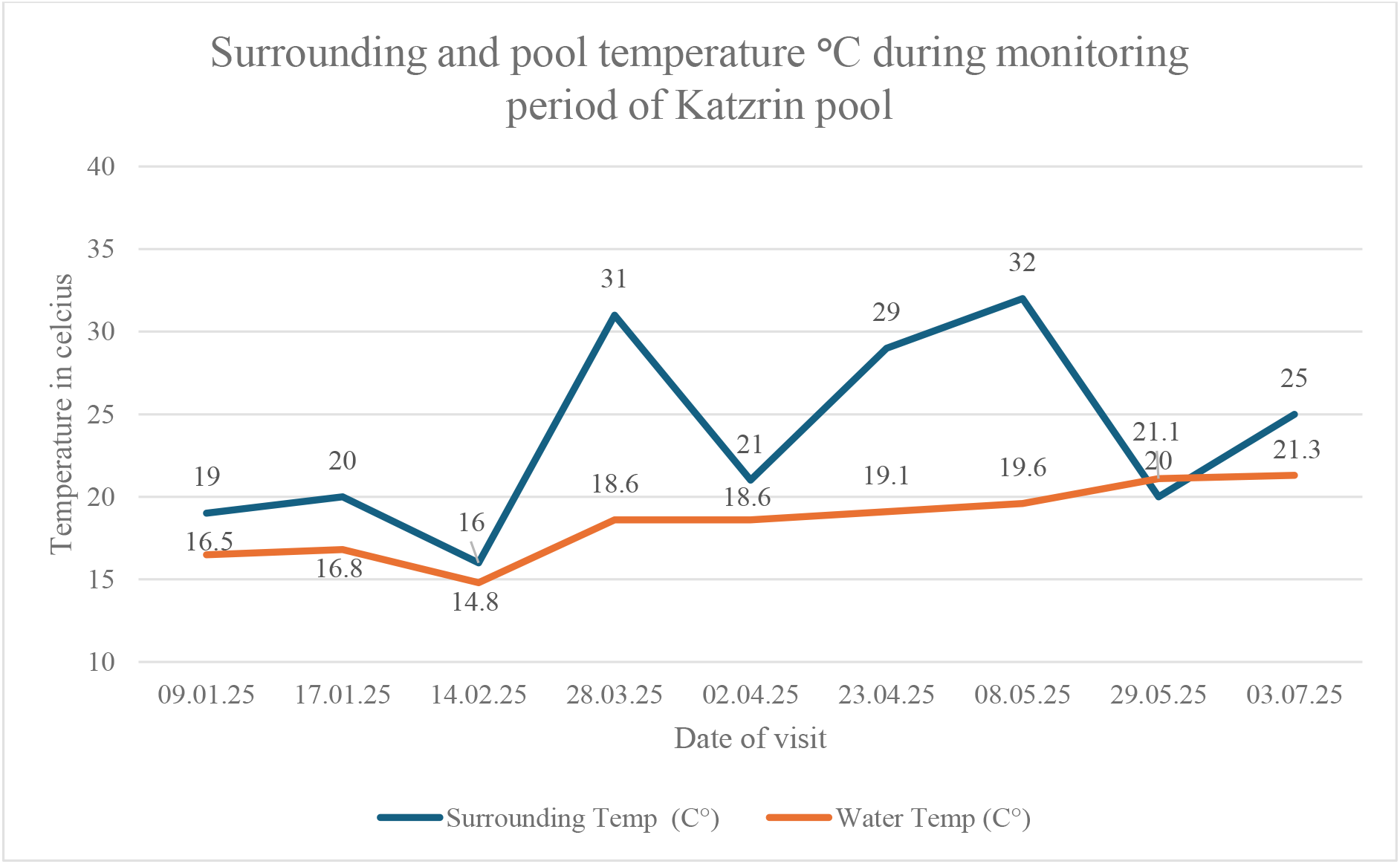
Seasonal variation in water temperature (°C) recorded at Zavitan Stream Katzrin Pool between January and July 2025.

**Figure 4.**
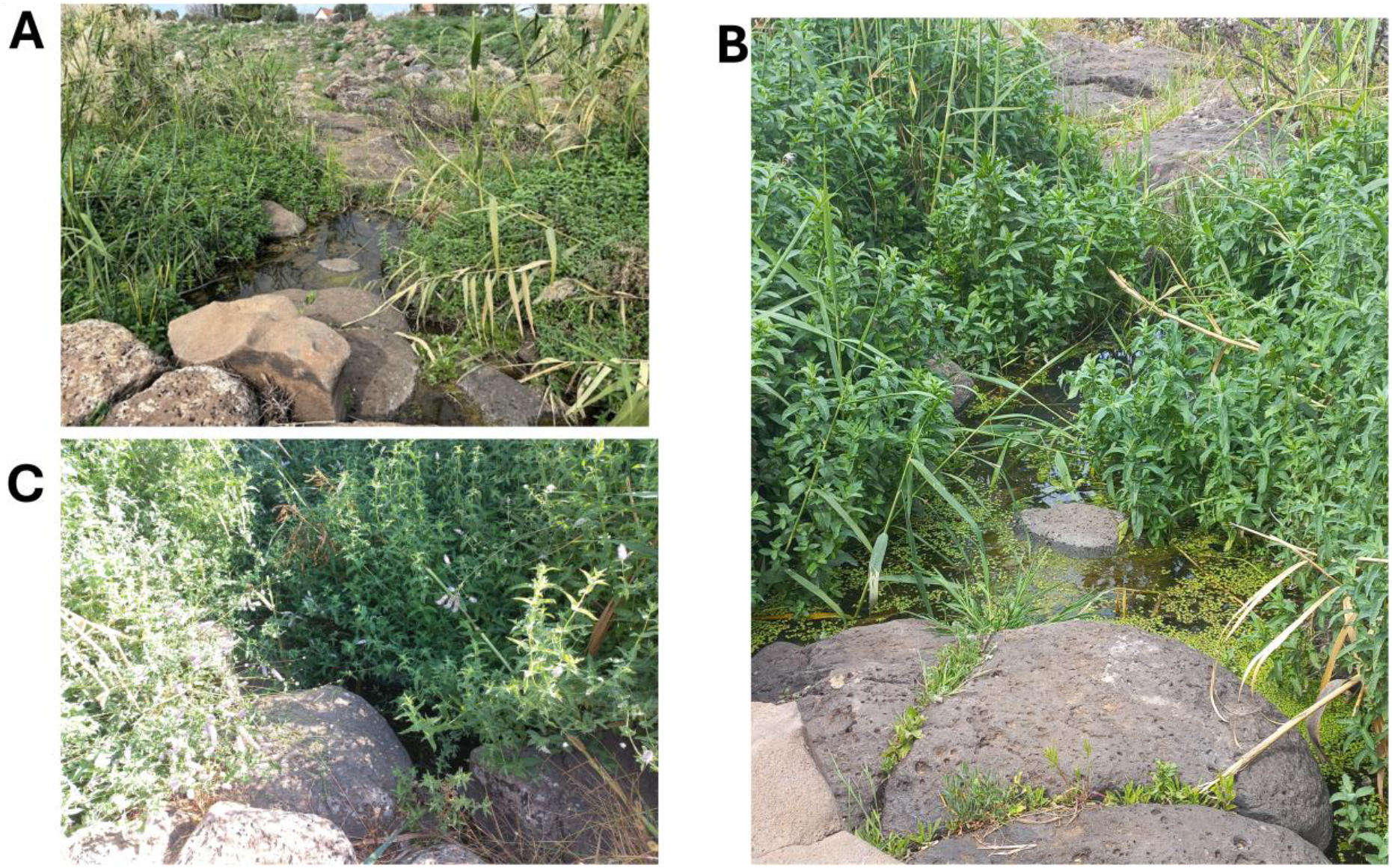
Seasonal changes in the hydrological and vegetative condition of Zavitan Stream Katzrin Pool during the 2025 monitoring period. (A) The pool at the beginning of the monitoring period in January, showing a high-water level and sparse vegetation. (B) The pool in March, characterized by increased aquatic and marginal plant growth. (C) The pool during the final visit in July, when only minimal residual water remained prior to complete desiccation.

Ambient air temperature measured during field visits ranged from 16 °C to 32 °C, with a mean of 23.67 °C. The highest air temperatures were recorded during late spring and early summer visits, coinciding with accelerated evaporation and declining water levels.

Overall, the abiotic conditions reflected a typical Mediterranean seasonal hydroperiod pattern, characterized by winter inundation, progressive spring warming, and complete summer desiccation.

### Animal Species in Zavitan–Katzrin Pool

Anuran larvae (tadpoles) and small juvenile frogs were observed during multiple sampling visits, indicating active amphibian reproduction within the pool. On one sampling occasion, three Eppendorf tubes containing whole larvae were collected for documentation and taxonomic identification.

Throughout the hydroperiod, the pool supported a diverse aquatic community. Observations included abundant anuran larvae (frogs and toads), Odonata nymphs (dragonflies and damselflies), aquatic hemipterans (water bugs), and small freshwater gastropods, including *Melanopsis praemorsa*.

The presence of multiple trophic groups, comprising both invertebrate and vertebrate taxa, indicates that the pool functioned as an ecologically active seasonal wetland habitat during the monitoring period.

### Morphological Description of the Amphibian Specimen

Our observations of the local amphibian species inhabiting Katzrin seasonal pool revealed an anuran species was present, most likely sp. Levant water frog (*Pelophylax bedriagae*). This amphibian is typical to the streams, permanent pools, and winter ponds in the Golan Height and galilee region.

The adult Levant water frog (*Pelophylax bedriagae*) is characterized by an olive to bright green dorsal coloration bearing irregular dark spots or blotches. In many individuals, a pale vertebral stripe extends along the midline of the dorsum, although this stripe may be absent in some specimens. The tympanum (eardrum) is relatively large and clearly visible posterior to the eye.

Figure 5. (upper left) displays the adult form and the late-stage metamorphosing (right side) with characteristics typical of the final phases of larval development prior to complete tail resorption. The specimens exhibited relatively robust and compact body, with a broad head and well-developed trunk musculature. Both hind limbs are fully formed, elongated, and muscular, bearing distinct digits adapted for terrestrial locomotion. The forelimbs were also completely emerged, indicating that the individual has reached the metamorphic climax stage.

**Figure 5.**
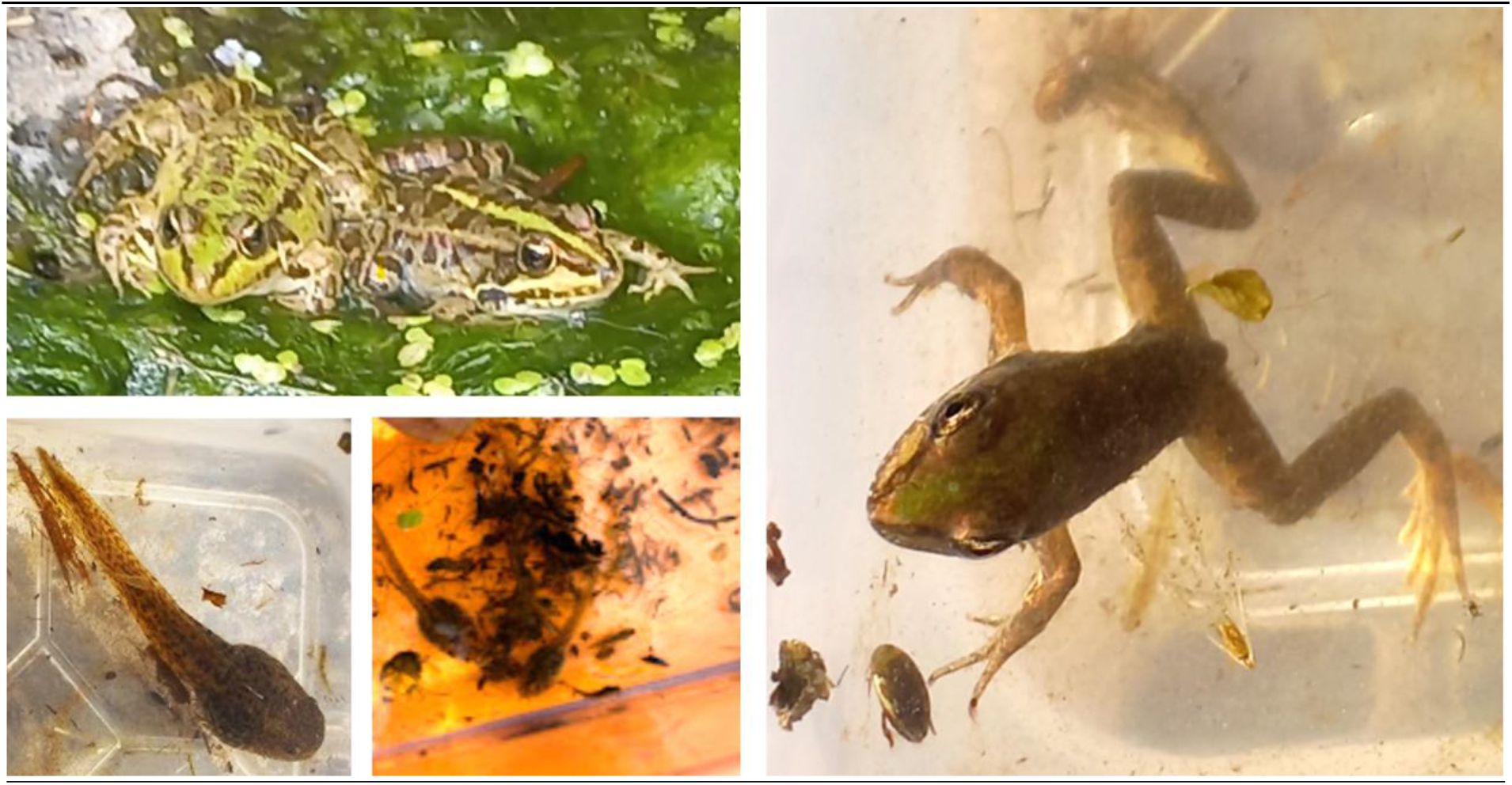
Morphological Description in Zvitan-Katzrin Pool. Image of Levant Water Frog (*Pelophylax bedriagae)*.

**Figure 6.**
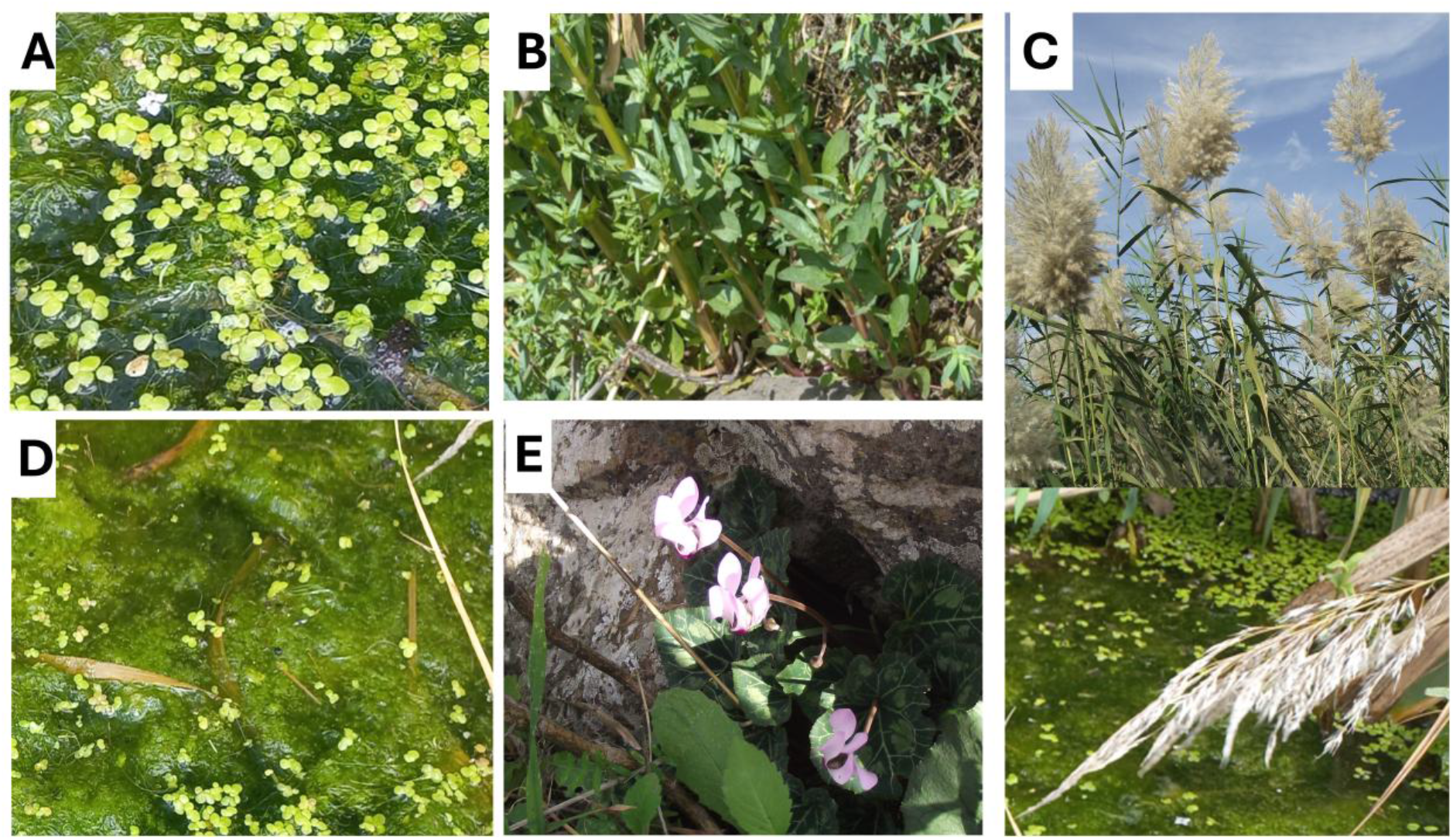
Representative plant species observed at Zvitan–Katzrin Pool during the 2025 hydroperiod. (A) Duckweed (*Lemna spp*.). (B) Water mint (*Mentha aquatica*). (C) Common reed (*Phragmites australis*). (D) Filamentous green algae (unidentified taxa). (E) Persian cyclamen (*Cyclamen persicum*) in the surrounding terrestrial habitat.

Figure 5. (Both images at the lower left) displays tadpoles which are thought to be of the species Levant water frog. The tadpoles of this species are described as oval shaped with dark brown to black body, long muscular tail with transparent fin membranes and dorsally positioned eyes. It is adapted to permanent or semi-permanent freshwater habitats and feeds mainly on algae and detritus. Its metamorphosis typically occurs in late spring to summer, depending on water temperature and hydroperiod.

During the 2025 hydroperiod, the Zvitan–Katzrin Pool supported several representative aquatic and riparian plant species. Floating mats of duckweed (*Lemna* spp.) covered parts of the water surface. Water mint (*Mentha aquatica*) grew along the shallow margins of the pool, while common reed (*Phragmites australis*) formed dense stands in the littoral zone. Within the shallow water column, filamentous green algae developed in sun-exposed areas. In the surrounding terrestrial habitat, Persian cyclamen (*Cyclamen persicum*) was recorded among the Mediterranean vegetation. These plant assemblages contributed to the structural complexity of the temporary freshwater habitat and likely provide microhabitats for aquatic invertebrates and amphibians.

## Discussion

The present study provides a focused ecological assessment of a seasonal pool located within the Zavitan (Zvitan) Stream drainage near Katzrin in the central Golan Heights. The results indicate that this pool functions as a typical Mediterranean temporary wetland, characterized by winter inundation, gradual warming during spring, and complete desiccation by midsummer. Such seasonal hydrological dynamics are characteristic of Mediterranean temporary freshwater systems, where precipitation during winter leads to the formation of ephemeral aquatic habitats that gradually dry during the warm and dry summer months (Gasith and Resh, 1999; Williams, 2006). Similar hydrological regimes have been documented in basaltic landscapes of northern Israel and the Golan Heights, where winter rainfall and impermeable basaltic substrates promote the formation of temporary pools and streamside wetlands (Gvirtzman, 2009; Sneh and Weinberger, 2003). These environmental conditions play an important ecological role in supporting amphibian breeding activity and diverse assemblages of aquatic invertebrates typical of Mediterranean temporary waters (Gasith and Resh, 1999; Yom-Tov and Tchernov, 1988).

The hydroperiod extended from January until July 2025, indicating a relatively prolonged water retention phase compared to shallow ephemeral depressions in more arid zones of northern Israel (Degani and Kaplan, 1999). Water temperature ranged between 14.8 °C and 21.3 °C, with gradual seasonal warming. Such temperature ranges fall within the optimal developmental window for most Mediterranean anuran larvae, facilitating rapid growth prior to desiccation (Degani, 2024a). The progressive increase in ambient temperature (up to 32 °C) likely accelerated evaporation and contributed to the contraction of aquatic habitat during late spring. In Mediterranean temporary systems, shortening hydroperiods are known to impose strong selective pressures on larval growth rate, metamorphic timing, and survival (Goldberg et al., 2009; Degani, 2024a). The complete desiccation observed in July confirms that successful recruitment depends on the synchronization between breeding phenology and hydroperiod length. The amphibian community structure previously described for the Golan Heights by Degani and Kaplan (1999) allowed comparison between earlier observations and the results obtained in the present study. Some species previously reported from this area were not detected during the present survey. In particular, the banded newt (Ommatotriton vittatus), which had been recorded in the region, was not observed in the studied pool. This absence may reflect several limiting ecological factors. Firstly, the pool posed a Hydroperiod limitation. While the lentic environment persisted until July, the site may lack the requisite depth stability necessary to support the prolonged larval development characteristic of urodeles. Secondly, Banded newts often prefer shaded, spring-fed, or semi-permanent pools within canyon systems, a profile that contrasts with the more exposed nature of the current site. (Degani and Ahkked, 2021). The Zvitan–Katzrin pool represents an intermediate ecological unit within the broader Zavitan drainage system. While canyon sections of the stream may maintain semi-permanent water through spring discharge, plateau pools such as the one studied here function as seasonal breeding sites primarily for anurans.

The absence of urodeles and the dominance of anuran larvae reinforce the importance of hydroperiod duration and habitat structure in shaping amphibian assemblages in the central Golan Heights. Temporary pools serve as critical reproductive habitats for many amphibian species but are highly sensitive to interannual climatic variability. These ephemeral wetlands typically depend on winter precipitation and seasonal hydrological cycles to maintain water availability during the larval development period (Gasith and Resh, 1999; Williams, 2006). Even modest reductions in winter rainfall or increases in spring temperature can substantially shorten the hydroperiod of temporary pools, thereby reducing the time available for larval growth and metamorphosis and consequently decreasing larval survival probability (Semlitsch, 2003; Wellborn et al., 1996). Such hydrological instability is a characteristic feature of Mediterranean climate regions, where amphibian reproductive success is closely linked to the duration and persistence of temporary aquatic habitats (Gasith and Resh, 1999).

## Conclusion

The Zavitan–Katzrin seasonal pool functions as an ecologically active temporary wetland supporting diverse aquatic invertebrates and successful anuran reproduction during its hydroperiod. The absence of banded newts suggests habitat-specific constraints or distributional fragmentation within the central Golan Heights.

Hydrological regime, temperature patterns, and vegetation structure collectively determine amphibian assemblages in this system. Given the sensitivity of Mediterranean temporary wetlands to climatic variability, the studied pool provides a valuable baseline for future ecological monitoring and conservation planning within the Zavitan Stream basin.

## Acknowledgements

The authors thank Yehonatan Almozlino and Avner Man for their assistance and collaboration during ecological fieldwork.

## Author Contributions

Monika Almozlino - Writing – review & editing, Writing – original draft, Data curation, Formal analysis, Investigation, Visualization, Project administration, Formal analysis.

Gad Degani - Writing – review & editing, Supervision, Conceptualization, Project administration, Investigation, Resources, Funding acquisition.

Dani Bercovich - Writing – review & editing, Supervision, Funding acquisition, Project administration.

Ari Meerson - Writing – review & editing, Supervision.

The authors report there are no competing interests to declare.

## References

Cox, C.B., Moore, P.D., 2001. Biogeography: an ecological and evolutionary approach. 6th ed. Blackwell Science, Oxford.

Danin, A., 2004. Distribution Atlas of Plants in the Flora Palaestina Area. Israel Academy of Sciences and Humanities, Jerusalem.

Degani, G., Kaplan, D., 1999. Distribution of amphibian larvae in Israeli habitats with changeable water availability. Hydrobiologia 405, 49–56.

Degani, G., 2019. The Fire salamandra (Salamandra infraimmaculata) and the Banded newt (Triturus vittatus) along the southern border of their. Published by Scientific Research Publishing Inc. ISBN, 978-971-61896-61693-61893.

Degani, G., 2024a. Biological Adaptations of Anuran Species across Diverse Habitats, Spanning Mediterranean to Desert Climates. Scientific Research Publishing, 1–92.

Degani, G., 2024b. Differences in Ecological and Genetic Adaptations between Salamandra infraimmaculata and Ommatotriton vittatus. Open Journal of Animal Sciences 14.

Degani, G., Ahkked, N., 2021. Ecological and Biological Adaptations of Triturus vittatus vittatus (Urodela) to an Unstable Habitat. Int. J. Zoo. Animal. Biol. 3, 1–8.

Degani, G., Goldberg, T., 2013 Aquatic Invertebrates in Different Bodies of Water in a Semi-Arid Zone American Open Animal Science Journal 1, 1–15.

Gafny, S., Goren, M., Gasith, A., 2000. Habitat condition and fish assemblage structure in a coastal Mediterranean stream (Yarqon, Israel) receiving domestic effluent. Hydrobiologia 422/423, 319–330.

Gasith, A., Resh, V.H., 1999. Streams in Mediterranean climate regions: abiotic influences and biotic responses to predictable seasonal events. Annual Review of Ecology and Systematics 30, 51–81. 10.1146/annurev.ecolsys.30.1.51

Goldberg, T., Nevo, E., Degani, G., 2009. Breeding site selection according to suitability for amphibian larval growth under various ecological conditions in the semi-arid zone of northern Israel. Ecologia Mediterranea 35, 65–74.

Goldberg, T., Nevo, E., Degani, G., 2012. Amphibian Larval in Various Water Bodies in the Semi-arid Zone. Zoological Studies 51 345–361.

Gvirtzman, H., 2009. Israel Water Resources: From the Prehistoric Era to the Present. Yad Ben-Zvi Press, Jerusalem.

Pearlson, O., Degani, G., 2007. Triturus v. vittatus (Urodela) larvae at various breeding sites in Israel. Progrese şi Perspective in Medicina Veterinară-Lucrări ştiinţifice 50, 214–226.

Por, F.D., 1975. An outline of the zoogeography of the Levant. Zoological Journal of the Linnean Society 57, 37–51.

Preißler, K., Küpfer, E., Löffler, F., Hinckley, A., Blaustein, L., Steinfartz, S., 2021. Genetic diversity and gene flow decline with elevation in the Near Eastern fire salamander a(Salamandra infraimmaculata) at Mount Hermon, Golan Heights. Amphibia-Reptilia 42, 241–247.

Roll, U., Dayan, T., Simberloff, D., 2009. Non-indigenous terrestrial vertebrates in Israel and their potential impacts on biodiversity. Biological Invasions 11, 1863–1875.

Semlitsch, R.D., 2003.Amphibian Conservation. Smithsonian Institution Press, Washington, DC.

Sneh, A., Weinberger, R., 2003. Geological map of Israel. Geological Survey of Israel, Jerusalem.

Udvardy, M.D.F., 1975. A classification of the biogeographical provinces of the world. IUCN Occasional Paper 18, International Union for Conservation of Nature and Natural Resources, Morges, Switzerland, 48 pp.

Wellborn, G.A., Skelly, D.K., Werner, E.E., 1996. Mechanisms creating community structure across a freshwater habitat gradient. Annual Review of Ecology and Systematics 27, 337–363.10.1146/annurev.ecolsys.27.1.337

Williams, D.D., 2006. The Biology of Temporary Waters. Oxford University Press, Oxford.

Yom-Tov, Y., Tchernov, E., 1988. The zoogeography of Israel: the distribution and abundance at a zoogeographical crossroad. Monographiae Biologicae 62, Dr. W. Junk Publishers, Dordrecht, 600 pp.

Zohary, D., Ostrovsky, I., 2011. Introduction to the flora of Israel. Israel Academy of Sciences and Humanities, Jerusalem.

